# Limitations of DNA barcoding in determining the origin of smuggled seahorses and pipefishes

**DOI:** 10.1101/2020.12.09.417998

**Authors:** Conny P. Serite, Ofentse K. Ntshudisane, Eugene Swart, Luisa Simbine, Graça L. M. Jaime, Peter R. Teske

## Abstract

Seahorses and pipefishes are heavily exploited for use in Traditional Chinese Medicine (TCM), and less frequently for curio markets or as aquarium fish. A number of recent studies have used DNA barcoding to identify species sold at TCM markets in East Asia, but the usefulness of this approach in determining the region of origin remains poorly explored. Here, we generated DNA barcodes of dried seahorses and pipefishes destined for TCM that were confiscated at South Africa’s largest airport because they lacked the export permits required for the CITES-listed seahorses. These were compared with published sequences and new sequences generated for Mozambican seahorses, with the aim of determining whether it is possible to identify their country of origin. All pipefishes were identified as *Syngnathoides biaculeatus*, a widespread Indo-Pacific species, but the published sequence data did not provide sufficient resolution to identify the region of origin. The same was true of the majority of seahorses, which could not even be identified to species level because they clustered among an unresolved species complex whose sequences were published under the names *Hippocampus kuda, H. fuscus* and *H. capensis*. The presence of a few specimens of a second seahorse, *H. camelopardalis*, suggests that the shipment originated from East Africa because the range of this seahorse is centred around this region, but again, it was not possible to determine their country of origin. Even though seahorses and pipefishes have high levels of genetic population structure because of their low dispersal potential, DNA barcoding was only suitable to tentatively identify species, but not their region of origin. DNA barcoding is increasingly used to identify illegally traded wildlife, but our results show that more sophisticated methods are needed to monitor and police the trade in seahorses and pipefishes.

## Introduction

The family Syngnathidae consists of over 300 species of fish. Seahorses and pipefishes belong to this family, along with pipehorses and seadragons [1–5]. Many syngnathids live in highly vulnerable inshore marine habitats [6] such as shallow reefs and lagoons [7,8], mangroves [9], estuaries [7,10,11], seagrass beds and algal flats [12,13]. This makes them highly susceptible to overfishing and habitat destruction. In addition, although some syngnathids can have thousands of young, brood sizes can be as small as 20 eggs or less per brooding male, making some species vulnerable to local extinction [10,14–16]. Some syngnathids may also form monogamous breeding pairs, and when one partner dies or is captured, the other stops reproducing until it finds another breeding partner [10,14,16,17], and declining population numbers may decrease the chances of finding a new mate.

In many regions, syngnathids are threatened by commercial overexploitation. Some are sold as curious or aquarium fish [2,5,8,9,14,18], but the Traditional Chinese Medicine (TCM) market is the main consumer of wild syngnathids [12,14,18]. For example, the global annual seahorse trade exceeds 20 million individuals [19–22]. Dried syngnathids are used to treat a wide range of conditions such as impotence and infertility, asthma, kidney disorders and other bodily illnesses [2,12,23,24].

All seahorse species (*Hippocampus* spp.) are listed in Appendix II of the Convention on International Trade in Endangered Species of Wild Fauna and Flora (CITES). In contrast, pipefishes (several genera) have not been afforded the same status, and most are listed as of ‘least concern’ or ‘data deficient’, even though they often face the same threats as seahorses. Only one species, *Syngnathus watermeyeri* Smith, 1963, is listed as ‘critically endangered’ [2,14,25,26].

Although seahorses and pipefishes are readily identifiable at higher taxonomic levels, identifying species is notoriously difficult. Roughly 42 seahorse species [27] and about 200 pipefish species occur worldwide [28], but the exact number of species that exist is contested. For example, between 41 [6] and 57 [29] seahorses are currently considered valid by different authors.

A combination of morphological characteristics [6,30] and genetic methods [3,5,31] holds great promise in improving this situation. In particular, DNA barcoding (which involves the sequencing of a portion of the mitochondrial DNA COI gene) tends to have sufficient genetic variation to distinguish closely related species from each other [32–34]. This genetic marker was successfully used by Hou et al. [31], Zeng et al. [5] and Wang et al. [35] to identify dried seahorse and pipefish samples to species level. In the present study, we used this marker to determine whether it provides enough resolution to determine the country of origin of illegally traded syngnathids confiscated at South Africa’s largest airport. Identifying the country of origin of such samples is important to uncovering illegal smuggling operations and policing them at their source.

## Materials and methods

### Samples

The majority of the samples used in this study originated from outbound mail parcels containing dried seahorses, pipefishes and sea cucumbers that were confiscated at OR Tambo International Airport in Johannesburg, South Africa. This airport, the busiest in Africa, is one of several locations where such seizures are made on a regular basis (others being the Libombo border crossing with Mozambique, and the Chinese market in the Cyrildene/Bruma area in Johannesburg). The seahorses (which are CITES listed) were not accompanied by any documentation, and it could not be ruled out that some of them were Knysna seahorses (*Hippocampus capensis*), an Endangered species that is endemic to South Africa [36].

Of the large number of morphologically similar dried samples intercepted by Biodiversity Enforcement, we selected a sample of 11 seahorses and 8 pipefishes. Moreover, given the scarcity of Mozambican samples in the genetic databases, seven additional Mozambican seahorses from a curio shop in Pemba that were morphologically similar to the confiscated specimens were also processed. To avoid supporting this trade, only fin or tail clippings were used, and no dried seahorses were purchased. Examples of the dried syngnathids confiscated at OR Tambo are shown in Fig. 1.

**Figure 1.**
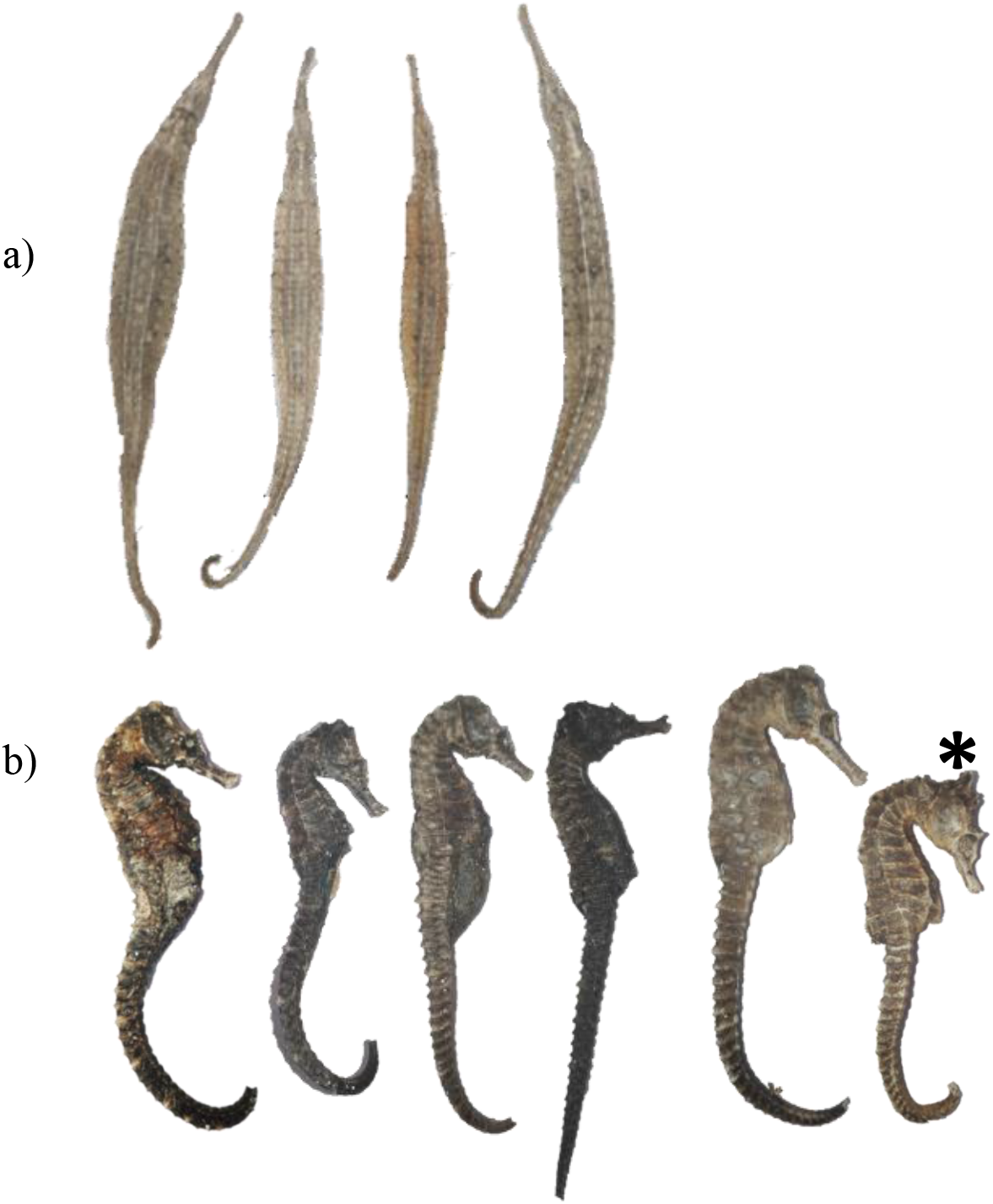
Examples of (a) pipefish and (b) seahorse samples confiscated at OR Tambo International Airport that were processed in this study. The seahorse marked with an asterisk (*) was identified as *Hippocampus Camelopardalis*.

### DNA extraction, amplification and sequencing

Mitochondrial DNA was extracted using the CTAB protocol [37]. Small pieces of tissue were cut from each specimen using a sterile scalpel, and the tissue samples were then placed into 1.5 ml Eppendorf tubes. One ml of CTAB buffer (2 g of CTAB powder, 1 g of PVP powder, 10 ml of 1 M Tris, 28 ml of 5 M NaCl, 4 ml of 0.5 M EDTA and filled up to 100 ml with ddH_2_O) and 10 μl of Proteinase K were then added to the tubes. The tubes were kept at 55°C overnight and vortexed occasionally until the tissue samples had completely dissolved. Then, 500 μl of a 1:24 isoamyl alcohol-chloroform mixture was added to the tubes, which were shaken vigorously for 30 s until the mixture had turned completely white. The tubes were then centrifuged at 13 000 rpm for 5 min until two layers of liquid formed, separated by a layer of foam. Up to 500 μl of the clear top liquid layer was transferred to new 1.5 ml Eppendorf tubes, followed by addition of 500 μl of isopropanol. The tubes were then vortexed and centrifuged at 13 000 rpm for 10 min. Tubes were decanted, and the DNA pellets were washed twice using 700 μl of 70% ethanol, in each case followed by centrifugation at 13 000 rpm for 10 min. All the liquid was then removed from the tubes using a narrow bore pipette, and the DNA pellets were left to dry at room temperature for approximately 30 min. Lastly, 40 μl of dilute TE buffer (500 μl of 1 x Tris-EDTA buffer stock and 49.5 ml ddH_2_0) was added to each Eppendorf tube, and the tubes were vortexed and stored at −80 °C until further processing.

The DNA barcoding marker for animals (i.e., the mitochondrial cytochrome oxidase c subunit I gene, abbreviated as COI or cox1) was amplified using primers mlCOIintF and jgHCO2198 [38]. These primers produce short amplified fragments of ~313 bp and are thus more likely to amplify PCR products when DNA is degraded (as in the case of dried syngnathids) than the fragment amplified using the more commonly used primers by Folmer et al. (1994). PCR amplifications were done using a total volume of 33 μl, which contained 3 μl of sample DNA and 30 μl of a master mix that contained 3 μl of 10 μM of each primer, 0.3 μl of Taq polymerase (5 units ml^-1^), 3 μl of 25 mM MgCl_2_, 3 μl of 1 μM of each dNTP and 14.7 μl of dH_2_O). Amplification was conducted on a MultiGene OptiMax thermal cycler using the following conditions: a denaturing step at 96°C for 5 minutes, 40 cycles of 30 s at 96°C for denaturation, 2 min at 49°C for annealing and 50 s at 72°C for extension, followed by a final extension 72°C for 10 min. Successful amplification was confirmed using 2% agarose gel electrophoresis with 0.5X TBE buffer, and the bands were visualized using GelRed Nucleic Acid Gel Stain (Biotium). The PCR products were sequenced by the Central Analytical Facilities (CAF) at Stellenbosch University.

### Data analysis

To identify syngnathids that are closely related to the species processed in this study, we conducted both BLAST searches of GenBank entries [40] and, based on preliminary results, searched for matches on BOLD [41] based on taxonomic identifications. The resulting sequences were then aligned manually in MEGA X [42]. Phylogenetic trees of the aligned sequences were constructed in MEGA X using the neighbour-joining algorithm (NJ; [43]) and the maximum-likelihood (ML) method. The evolutionary distances in all NJ trees were computed using the Kimura 2-parameter model (K2P; [44]), and the ML trees were constructed using the most appropriate model identified using the Bayesian information criterion (BIC; [45]). Pairwise deletion was applied to all trees, and nodal support was based on 1000 bootstrap replications [46]. Both the NJ and ML trees produced congruent results, so only the NJ trees were reported, and the bootstrap values of the ML trees were added to some nodes.

## Results

### DNA extraction, amplification sequencing and identification

Despite their dried state, the COI gene of all pipefishes and seahorses confiscated at OR Tambo amplified successfully, but the amplification of only five of the Mozambican seahorses was successful. These sequences have been submitted to the GenBank repository (accession numbers XX-XX). All the confiscated pipefish samples were identified as *Syngnathoides biaculeatus* (Bloch, 1785). All sequences of the confiscated samples were identical and clustered among published sequences of specimens collected throughout the species’ range (Fig. 2), with the sequence of a Tanzanian pipefish being most similar. One of the confiscated seahorses was identified as *H. camelopardalis* Bianconi, 1854 (marked with an asterisk in Fig. 1b), and its sequence was identical to the published sequences of a Tanzanian specimen and two of the three new Mozambican specimens (Fig. 3). The remaining seahorse samples (10 samples from OR Tambo and two from Mozambique) were difficult to identify because they clustered among published sequences of *H. fuscus* Rüppell, 1838, *H. kuda* Bleeker, 1852, and *H. capensis* Boulenger, 1900 (Fig. 4). As these samples also showed considerable morphological variation (Fig. 1b), they were labelled *Hippocampus* sp.

**Figure 2.**
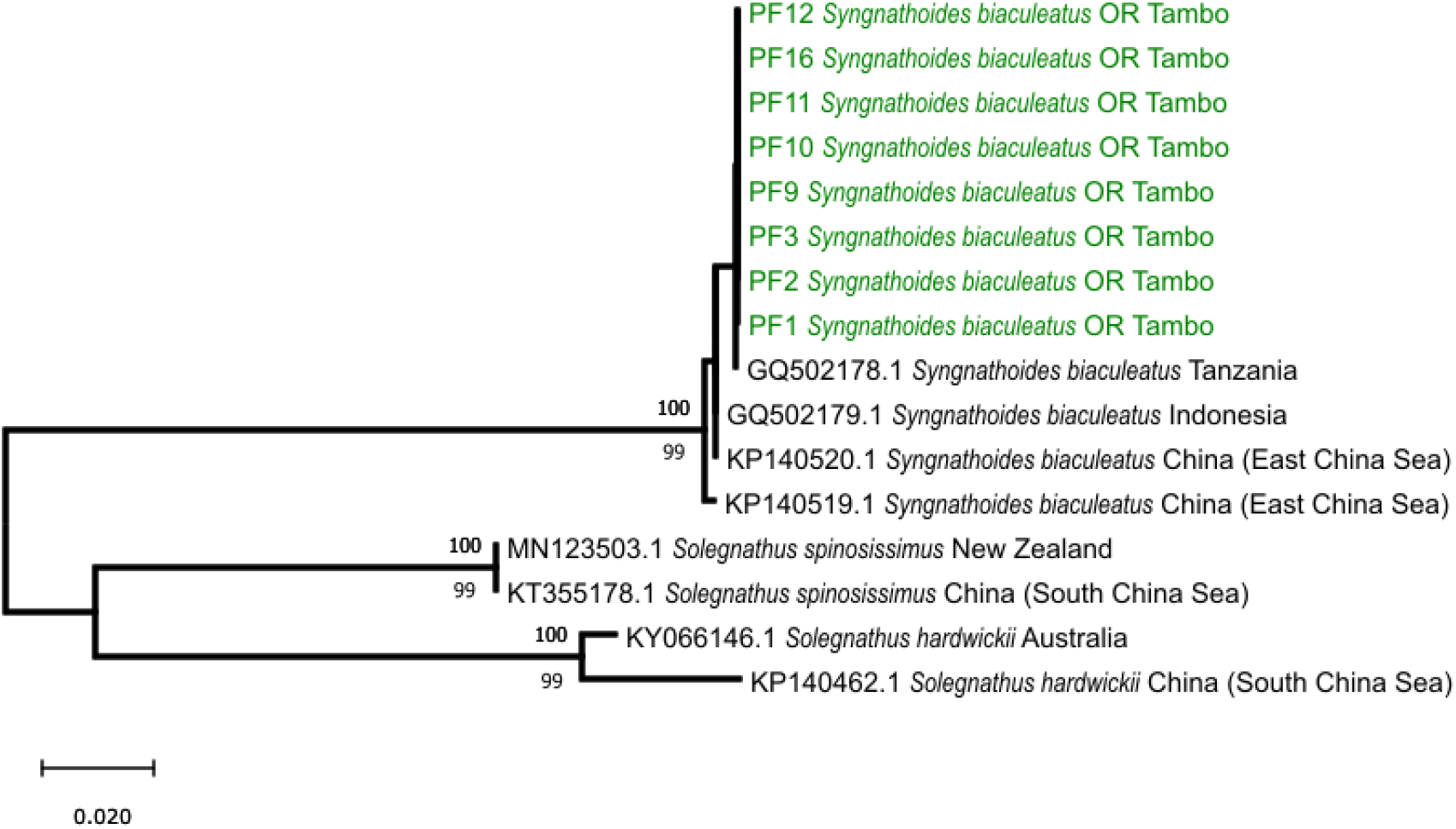
Neighbour-joining (NJ) phylogenetic tree constructed using COI sequences of *Syngnathoides biaculeatus* (green) from OR Tambo International Airport, and reference sequences of the same species from Tanzania, China and Indonesia. The outgroup included sequences of the closely related *Solegnathus spinosissimus* from New Zealand and China and *Solegnathus hardwickii* from Australia and China. Bootstrap values >75% for NJ and ML analyses are shown above and below some nodes, respectively. The HKY + G model [47] was used for the ML analyses.

**Figure 3.**
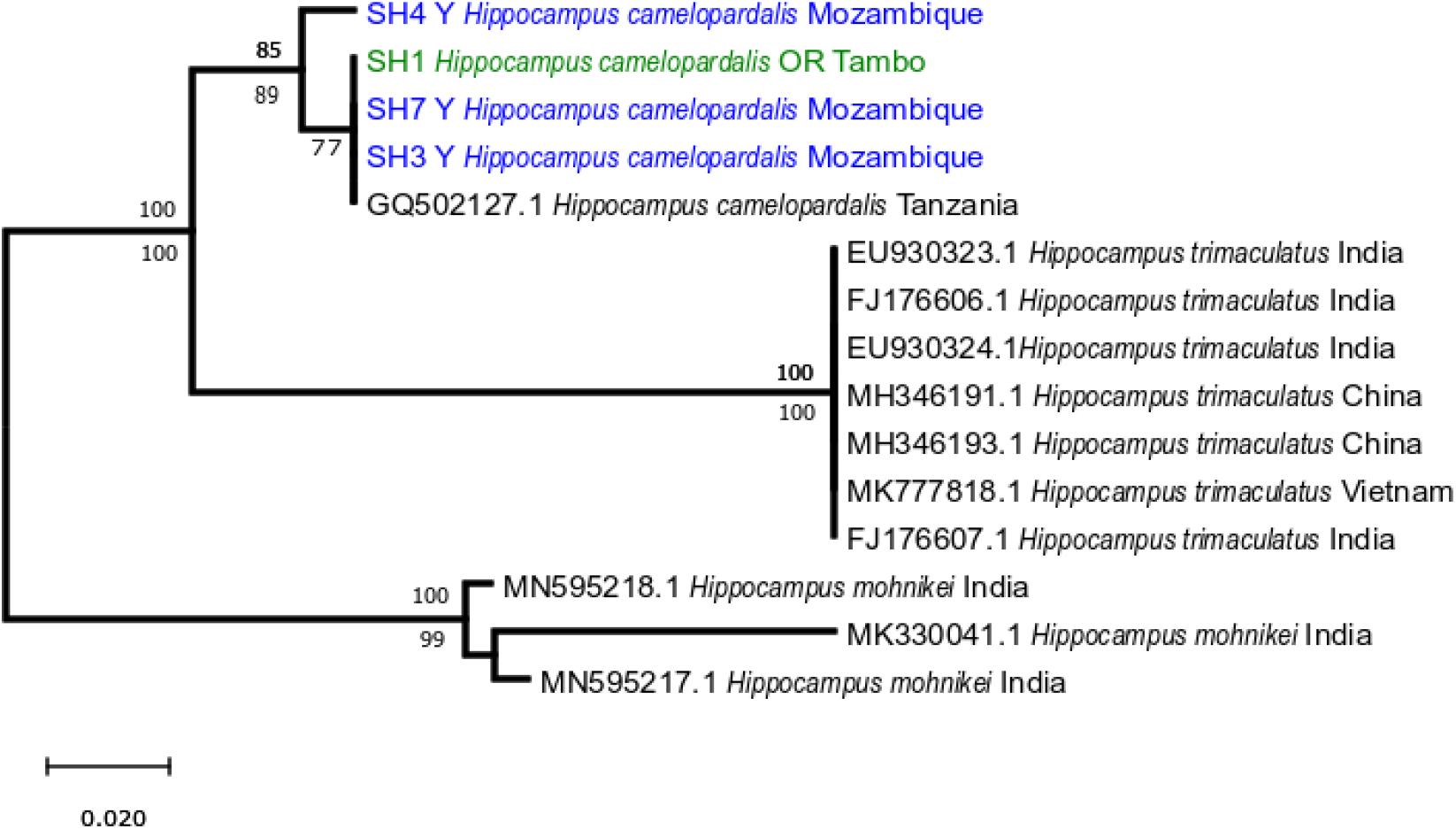
Neighbour-joining (NJ) phylogenetic tree constructed using COI sequences of *Hippocampus camelopardalis* from OR Tambo International Airport (green) and Mozambique (blue), and a reference sequence from Tanzania. The outgroup included published sequences of two closely related species, *H. trimaculatus* and *H. mohnikei*. Bootstrap values >75% for NJ and ML are shown above and below some nodes, respectively. The T92 model [48] model was used for the ML analyses.

**Figure 4.**
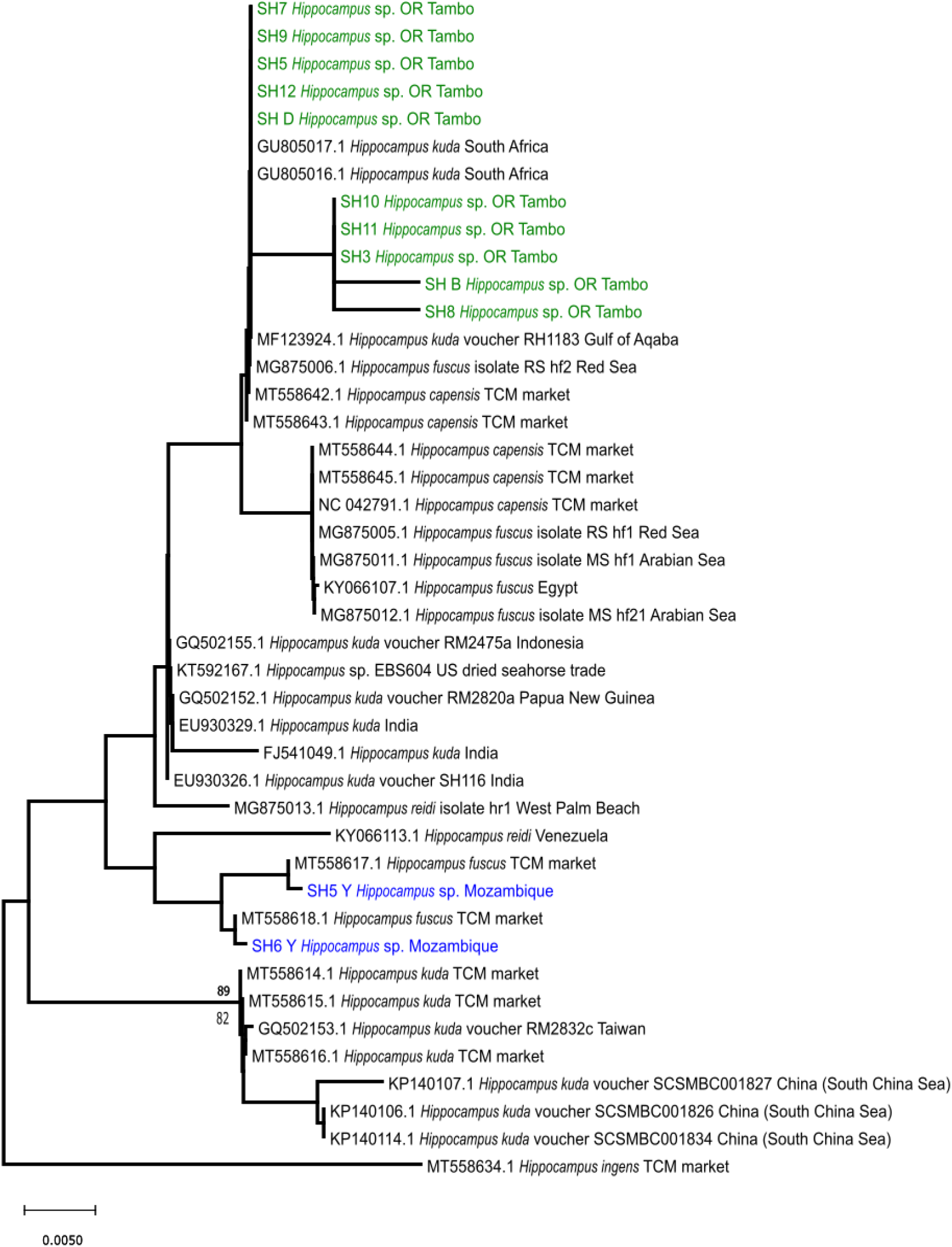
Neighbour-joining (NJ) phylogenetic tree constructed using COI sequences of *Hippocampus* sp. from OR Tambo International Airport (green) and Mozambique (blue), as closely related species from a variety of sources. Bootstrap values >75% for NJ and ML are shown above and below some nodes, respectively. The K2 + G model [44] was used for the ML analyses.

## Discussion

DNA barcoding is a useful tool for identifying illegally traded animal species that may be endangered or vulnerable, such as seahorses [22,31]. Even though the dried syngnathids used in this study had not been preserved under optimal conditions for genetic research, the COI gene was successfully amplified and sequenced in most samples. Three species were identified: the alligator pipefish, *Syngnathoides biaculeatus*, the giraffe seahorse, *Hippocampus camelopardalis*, and a second seahorse that remained unidentified. As mentioned below, these samples most likely originated from the Western Indian Ocean, suggesting that OR Tambo International Airport is an important transit location within a trans-national smuggling supply chain that supplies TCM markets with East African marine organisms.

*Syngnathoides biaculeatus* is widely distributed throughout the tropical Indo-Pacific, from the Red Sea and East Africa to Japan, Samoa, Australia and the Tonga Islands [12,14,16]. The presence of these pipefishes among confiscated samples of seahorses whose range is limited to the Western Indian Ocean (*H. camelopardalis*, see next paragraph), as well as their genetic similarity to the Tanzanian reference sequence, suggests that the pipefish samples sequenced here may have originated from East Africa. Despite its ‘Least Concern’ status in the IUCN Red List [49], populations of *S. biaculeatus* are undoubtedly under considerable pressure. The species has been used in TCM for more than 600 years [12] and is considered to be the most heavily exploited pipefish species, although comprehensive range-wide figures are lacking [12,14]. The annual trade of dried pipefish species into Taiwan was reported to be around 1 600-16 500 kg between 1983 and 1993 [12,14], and the annual import of a mixture of pipefish into Hong Kong during 1998-2002 was estimated to have been around 7 500-21 300 kg, with further trade suspected to be occurring in India, Malaysia, Thailand and the Philippines [2,12,14]. Given that the samples sequenced in this study may have originated from East Africa, details on the export from this region are of particular interest. However, compared to imports, the details on legally traded exports from Tanzania (the only country from which such numbers are available) are likely a very significant underestimate, with 145 kg being officially exported in 1989, and roughly 4000 individuals reported to be sold monthly during the mid-1990s [18]. It is very likely that the vast majority of these pipefishes are traded unofficially.

*Hippocampus camelopardalis* is a species that is found in the Western Indian Ocean, from South Africa to Tanzania, and potentially north into Kenya [8,18,50]. It is listed as ‘Data Deficient’ on the IUCN Red List [50]. A more conclusive assessment of the origin of the *H. camelopardalis* specimen from OR Tambo is not possible because sequences have only been generated for specimens from Tanzania and Mozambique, and all formed a single genetic cluster. However, the presence of this species amongst the confiscated samples provides the most conclusive evidence that the smuggled syngnathids sequenced here originated from East Africa.

The seahorses referred to in this study as *Hippocampus* sp. were the most challenging to identify. Not only were they morphologically heterogeneous (Fig. 1b), but in this case, the genetic data were not even suitable to identify to which species they belong. The main reason for this is that the taxonomy of this group is unresolved. They are typically assigned to the *‘Hippocampus kuda*-clade’, which consist of various seahorse species, including *H. kuda, H. fuscus, H. borboniensis* Duméril, 1870, *H. capensis, H. algiricus* Kaup, 1856, and *H. reidi* Ginsburg, 1933 [6,11,21]. Teske et al. [11] referred to the East African members of this species as a unique Indian Ocean lineage of *H. kuda* that was distinct from a Pacific *H. kuda* lineage, and only seahorses from the Red Sea were referred to as *H. fuscus* on the grounds of being morphologically distinct. In a recent taxonomic revision, *H. fuscus* and *H. borboniensis* were considered to be synonyms of *H. kuda* due to the lack of distinguishable characters [6]. As a result of this confusion, published sequences of seahorses that may all be part of the same species were submitted under different names. Whereas *H. kuda* and *H. fuscus* are both acceptable names to describe these seahorses, the specimens referred to as “*H. capensis* TCM market” in Fig. 4 were misidentified, as this endangered species is endemic to South Africa. The mitochondrial genome that was used to identify these samples [51] was most likely from a specimen native to China that in other recent literature has been referred to as *H. casscsio* Zhang, Qin, Wang & Lin, 2016, and which may be a synonym of *H. fuscus* [52].

Overall, the trade in seahorses is somewhat better documented than the trade in pipefishes. Foster et al. [53] reported that the global annual trade of seahorses between 2004 and 2011 was approximately 3.3-7.6 million individuals, while Lawson et al. [54] estimated that the number could be much higher at approximately 37 million individuals. In 2019, 1.2 t of dried seahorses from Peru were seized in China, a shipment that was estimated to be worth 2.16 million US$ [55], and the considerable value of dried syngnathids thus becomes a tempting incentive for artisanal fishermen and smugglers. Among African countries, Tanzania is most important as a source for dried seahorses, with an estimated ~254 000 seahorses exported from to China in the year 2000 [8].

## Conclusion

Combined morphological and genetic identifications are useful in identifying dried seahorse and pipefish samples. However, even though such methods are already being used to identify illegally traded wildlife [56,57], the present study has highlighted several limitations. First, in order to identify species using DNA barcoding, it is important that a standardised reference database exists, as the majority of the seahorses sequenced in this study (i.e., those associated with the *Hippocampus kuda* complex) could not be identified to species level. Although a comparatively short fragment of the COI gene was sequenced here due to the low quality of the tissue samples, a longer fragment of this marker does not provide more taxonomic resolution [52]. However, a different mitochondrial marker (control region) was suitable to distinguish between *H. kuda* from the Indian Ocean, and a clade comprising *H. fuscus* and Chinese seahorses referred to as *H. casscsio* [52], suggesting that it may be possible to resolve the taxonomy of the *H. kuda* complex by increasing the number of genetic markers. Second, in cases where species could be identified, the genetic data were only suitable to identify major regions (e.g. the Indo-Pacific in the case of *S. biaculeatus*, and the Western Indian Ocean in the case of *H. camelopardalis*). Syngnathids tend to have low dispersal potential because they are poor swimmers [11,21,58], and studies using biparentally-inherited microsatellites have identified population structure at scales of 100 to 1000 km in some seahorses [36] and pipefishes [58]. To determine where extensive unregulated syngnathid fisheries exist that need to be monitored, genetic methods that are more informative than DNA barcoding are required that provide resolution at smaller geographic scales.

## Acknowledgements

We are grateful to Claudia Schnelle for assisting with the laboratory work. This study was financially supported by the University of Johannesburg (FRC/URC grant to PR Teske).

## Notes

### Competing Interest Statement

The authors have declared no competing interest.

